# Efficient learning and intrinsic noise filtering in recurrent spiking neural networks trained with e-prop

**DOI:** 10.1101/2025.05.25.656058

**Authors:** Davide Noè, Hideaki Yamamoto, Yuichi Katori, Shigeo Sato

**Affiliations:** Research Institute of Electrical Communication, Tohoku University, Japan; Graduate School of Engineering, Tohoku University, Japan; School of Systems Information Science, Future University Hakodate, Japan; International Research Center for Neurointelligence (WPI-IRCN), The University of Tokyo, Japan

**Keywords:** Recurrent neural networks, Spiking neurons, Eligibility propagation, Biologically plausible learning, Computational neuroscience

## Abstract

**Objective:** Biologically plausible learning rules for neural networks, such as e-prop (eligibility propagation), are essential both for advancing neuromorphic computing and for understanding fundamental mechanisms of learning in animal brains. However, their behavior under different network conditions remains unclear.

**Approach:** Here, we investigate the performance of the e-prop learning algorithm in recurrent spiking neural networks (RSNNs) across different levels of recurrent connectivity and input noise using a complex temporal credit assignment task, a supervised classification problem known to be solvable by rodents.

**Main results:** We show that increased sparsity in the recurrent layer significantly enhances learning performance by promoting the generation of more diverse activation patterns. Analysis of the network’s evolution further reveals that the e-prop-trained input layer evolves to route distinct inputs to different regions of the recurrent layer while suppressing the contribution of noise. This partially resembles signal routing functions attributed to the thalamus in mammalian sensory systems, providing additional support for the biological plausibility of e-prop.

**Significance:** These findings offer promising insights for efficiency and advantages of biologically inspired training in RSNNs.

## I. INTRODUCTION

With the burgeoning diffusion of machine learning [1, 2] in diverse fields such as health-care [3, 4], software engineering [5], and renewable energy research [6], the amount of data and power required to train and operate artificial neural network models is increasing rapidly [7]. This trend has renewed our attention to the initial inspiration for artificial intelligence: the brain [8, 9]. Indeed, biological brains can learn with limited data [10] and exhibit remarkable power efficiency. For example, OpenAI’s GPT-3 consumes roughly 64,350 MWh for training and further 0.3 Wh for each query [11], while the human brain continually operates on around 20 W [12]. These advantages sparked great interest in neuro-morphic engineering, leading to the development of brain-inspired hardware such as Intel’s Loihi [13] or our analog VLSI chip [14], which implement models that resemble biological brains: recurrent spiking neural networks (RSNN) [15, 16].

RSNNs have been the subject of studies throughout the years, but researchers have struggled to overcome the underlying intricacies of the training process, such as the non-linear nature of the spikes [17] or the large computational cost of unfolding the network in time to perform backpropagation (BPTT) [18]. Several approaches have been proposed to solve this issue, including spike timing dependent plasticity [19], neoHebbian three factor learning [20], and fast learning without synaptic plasticity [21]. One strong candidate attracting attention is a biologically plausible learning algorithm called eligibility propagation (e-prop) [22] that is based on the tag-and-capture hypothesis for synaptic adaptation [20]. The algorithm postulates that a chemical trace (e.g. Ca^2+^ ions) accumulates in the synapse in proportion to neural activity and that the network is then flooded by a neurotransmitter (e.g., dopamine or norepinephrine) that acts as a top-down learning signal [22]. Hence e-prop is naturally local (traces accumulate inside a neuron through activation alone) and online (information is propagated only forward in time), eliminating the need to unfold the network in time as in BPTT.

The combination of spiking nature of RSNNs and the local/on-line nature of e-prop gathered attention not only as a new biologically plausible machine learning but also for applications in neuromorphic hardware. For instance, the feasibility of implementing e-prop into neuromorphic hardware and the consequent advantages in terms of power consumption has been demonstrated through its implementation in SpiNNaker2 [23, 24] and in ASICs such as ReckOn [25]. However, in order to facilitate optimal hardware implementation, it is still imperative to analyze how network structures and input conditions affect learning performance and network dynamics. For instance, the neuronal connection probability in the brain is typically sparse, estimated at approximately 8–20% in the mammalian cortex [26, 27], and mimicking such characteristics may further reduce wiring costs and power consumption in neuromorphic hardware. Additionally, biological networks are inherently robust to noise, a desirable feature in real-world application of SNNs.

In this paper, we investigate the optimal configuration for e-prop by systematically analyzing the effects of network sparsity in the recurrent layer and evaluate the noise tolerance of the model through controlled noise injection. As a benchmark, an RSNN with spiking input and continuous output is employed and trained to solve a complex temporal credit assignment task. The e-prop algorithm is used to train the input and recurrent layers while a standard gradient descent training is used for the output layer. We find that sparser networks create more diverse activation patterns in response to different inputs, leading to a significant improvement in task performance. Furthermore, the network also shows strong resilience to injected noise, which is found to be achieved by adapting the input layer to enhance the signal component in the input channels. These findings bolster the biological plausibility claims of the e-prop algorithm and provide a framework for optimizing network connectivity for classification tasks.

## II. METHODS

### Neuron and network models

Our RSNN comprises *N*_rec_ spiking neurons with a 50/50 split of two different types,leaky integrate-and-fire neurons (LIF) and adaptive LIF neurons (ALIF) [22, 28]. In both neuron types, the behavior is governed by a real-valued quantity *v*_*i*_ ∈ ℝ that we refer to as a membrane potential, borrowing the term from the biological analogous, which evolves according to the following equation:

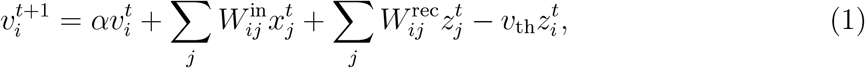

where 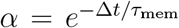 with *τ*_mem_ as the membrane time constant and Δ*t* as the time-step, *v*_th_ is the firing threshold of the neuron, 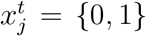 and 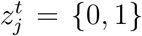 are respectively the input signal and the observable state (spike) of neuron *j* at time *t*. 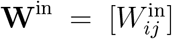 and 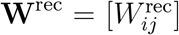 are synaptic weight matrices in the input and recurrent layers, respectively, and were initialized with a normalized Gaussian distribution 𝒩 (see Table I).

**Table 1:** List of parameters used in the experiments.

| Parameter | Value | Parameter | Value |
| --- | --- | --- | --- |
| $f_0$ | 4 Hz | $\Delta t$ | 1 ms |
| $v_{\text{th}}$ | 0.6 | $v_{\text{rest}}$ | 0 |
| $\tau_{\text{mem}}$ | 20 ms | $\tau_{\text{b}}$ | 2000 ms |
| $N_{\text{in}}$ | 40 | $N_{\text{rec}}$ | 100 (50 LIF + 50 ALIF) |
| $N_{\text{out}}$ | 2 | $W_{ij}^{\text{in}}$ | $\mathcal{N}(0, 1)/\sqrt{N_{\text{in}}}$ |
| $W_{ij}^{\text{rec}}$ | $\mathcal{N}(0, 1)/\sqrt{N_{\text{rec}}}$ | $W_{ij}^{\text{out}}$ | $\mathcal{N}(0, 1)/\sqrt{N_{\text{out}}}$ |
| $B_{ij}^{\text{out}}$ | $\mathcal{N}(0, 1)/\sqrt{N_{\text{out}}}$ | $\gamma_{\text{d}}$ | 0.3 |

Once *v*_*i*_ reaches a threshold value *v*_th_, a spike is generated according to:

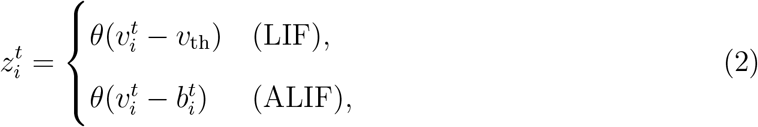

where *θ* is a Heaviside step function. For LIF neurons, *v*_*th*_ is a constant, while for ALIF neurons, the firing threshold *b*_*i*_ ∈ ℝ is defined as:

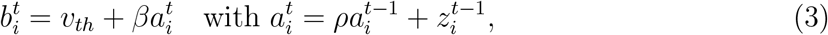

where 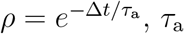 is the time constant for the adaptable threshold, and *β* is a constant. After a spike, *v*_*i*_ is reset in the subsequent time step using the last term in Eq. 1.

Finally, the output of the network 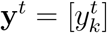 is given by:

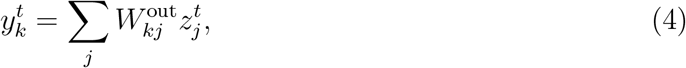

where 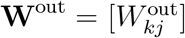 is the readout weight matrix, and *k* = (1, …, *K*) is the index for the classification category.

The three layers are initialized as a fully connected matrix, some synapses of the recurrent layer are then pruned to implement sparsity. These synapses are chosen differently in each iteration of the experiment but are kept fixed throughout each training process.

The values of each parameter are summarized in Table I.

### Task

In the complex temporal credit assignment task, an agent (a rodent in the original studies) is exposed to cues from left and right and needs to decide which side to take after a delay period [29, 30]. The input is given across *N*_in_ neurons as Poisson spikes [31] with frequency *f*_0_ divided among four channels as: left and right stimuli, the reward signal, and background noise.

Each episode comprises seven randomized input packets from left and right, followed by a delay period, and ends with the reward packet giving the correct answer to the agent. Each packet lasts for 150 ms, the delay period is 1050 ms thus the total length of an episode amounts to 2250 ms. This is a classification task with *K* = 2 categories, so we define the loss function *E* as the cross-entropy error in the following equation:

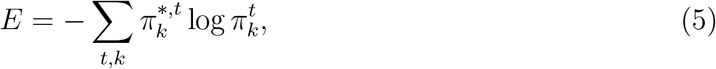

where 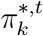 is a one-hot encoded *K*-dimensional target vector given as the reward packet, and 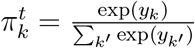.

During each epoch of training, the network is exposed to 64 randomly generated episodes. Testing is performed on 16 batches each containing 64 generated episodes. Unless otherwise noted, the noise-to-signal ratio was set to 50%, such that the probability of a spike being generated in the noise channel is half of the probability of generating a spike in a signal channel.

### Network evolution

We trained both **W**^in^ and **W**^rec^ using e-prop since all information the input and recurrent layers handle is in the form of spikes. The output signal, on the other hand, is continuous thus requiring a different learning algorithm for training **W**^out^; in this work, we employed a simple gradient descent method:

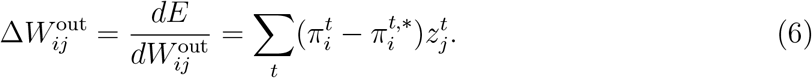

The e-prop algorithm is based on gradient descent:

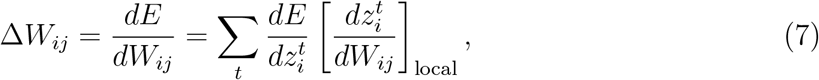

where 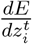 represents the learning signal *L*, which depends on the definition of the loss function. For the case at hand, *L* for neuron *i* at time *t* takes the form:

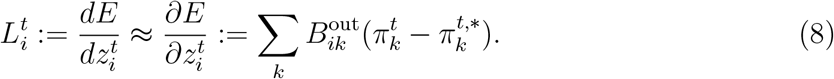

Here, we used an approximation from total to partial derivatives to preserve the online nature of the algorithm. In particular, using the total derivative would entail having information on the impact of 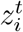 on *E* generated by future spikes of different neurons, which is not compatible with online information processing. This approximation involves substituting the output matrix **W**^out^ with a random fixed matrix 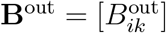, drawing inspiration from the broadcast alignment method [32].

The local term on the RHS of Eq. 7 is the eligibility trace (*e*_*ij*_), defined as:

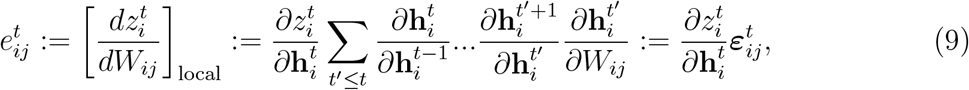

where 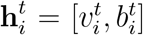 is the internal state of the network that evolves according to Eq. 1 and Eq. 3 to generate spiking activity in the visible state according to Eq. 2. The summed term on the RHS is labeled as the eligibility vector ***ε***_*ij*_ and can be computed recursively according to:

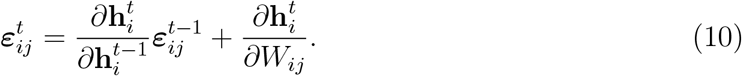

The spike function 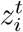 is non-differentiable, so its derivatives have been approximated by a tent function:

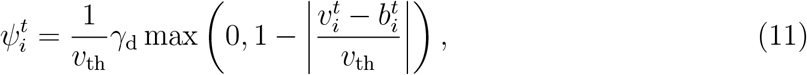

where *γ*_d_ is a constant. However, for the sake of clarity, we have written the derivatives of 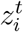 using the standard derivative symbol in a slight abuse of notation.

A more in-depth derivation of the traces for both the LIF and ALIF neurons can be found in the Supplementary Materials and in Ref. [22].

## III. RESULTS

Our RSNN model is represented in Fig. 1A. In this Section, we analyze the behavior of the model in solving the complex temporal credit assignment task (Fig. 1B). In particular, we studied the impact of connectivity on the e-prop learning algorithm as well as the network’s response to various amounts of input noise. We also analyzed the impact of learning on the landscape of synaptic connections. A schematic representation of the e-prop algorithm, along with an overview of the network’s dynamics and output before and after training, is presented in Figs. 1C and D.

**Figure 1:**
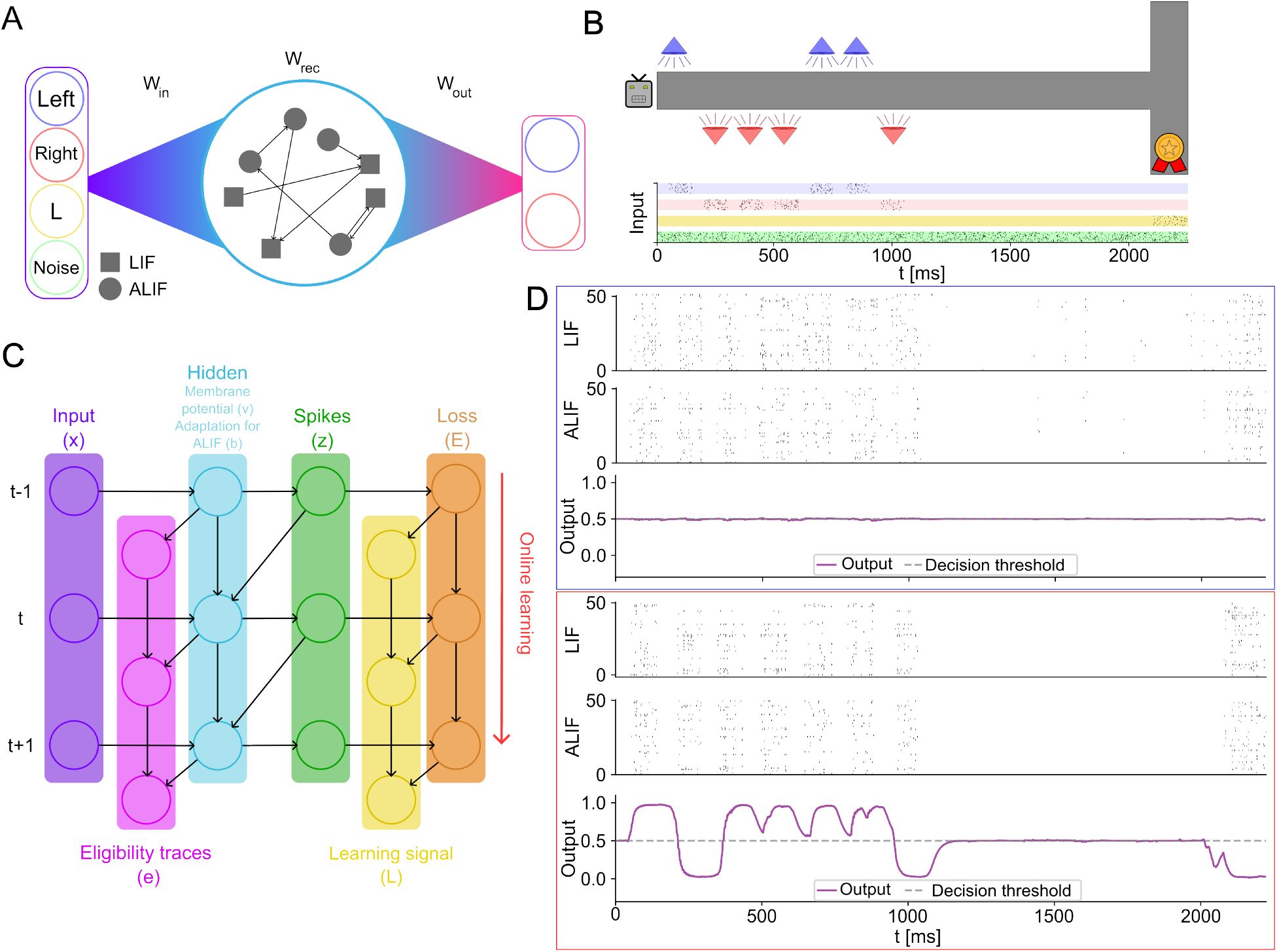
Overview of the experiment. A) Structure of the network. B) Visualization of the task along with the input signal for a single episode. The different input channels are highlighted using the following colors: left (blue), right (red), reward (yellow) and noise (green). C) Schematic representation of the e-prop learning algorithm. Black arrows represent the flow of information while time flows from top to bottom. D) Spiking activity and output during an an episode before training (top) and after the training with e-prop (bottom).

### A biologically realistic connectivity improves e-prop training performance

We first investigated how network connectivity impacts the behavior and performance of e-prop trained RSNNs. We focused on connection density, since neurons in the nervous system of animals are generally sparsely connected [26, 27, 33]. We started with a fully-connected network and pruned different randomly selected synapses (Fig. 2A). As summarized in Fig. 2B, networks with a lower connection density exhibited superior performance both in terms of loss after learning (loss row) and in terms of training efficiency (epoch row). Notably, we found that the optimal sparsity was at 92% (i.e., connection probability of 8%), showing the most efficient training in terms of epochs to achieve the accuracy threshold (set to 93% [22]). The average loss observed at that sparsity (0.18 ± 0.06, *n* = 20 samples) was not statistically different from the minimum average loss obtained at a sparsity of 99% (0.16 ± 0.03, *n* = 20 samples; two-sided *t*-test, *p >* 0.05). This connection density closely resembles that reported in murine studies [26], showing that a more biologically realistic network improves the performance of e-prop.

**Figure 2:**
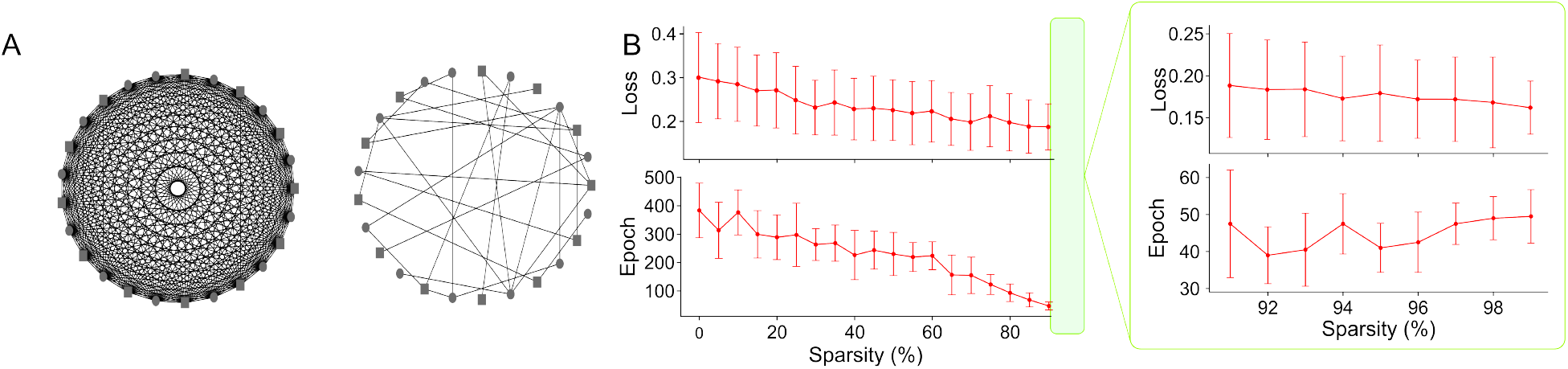
A) Fully connected network (left) compared to a network with optimal connectivity (92% sparsity; right). B) Model performance as a function of network sparsity. A zoom between 90% and 99% sparsity is shown on the right. The loss row presents the average loss between epochs 400 and 500. The epoch row presents the amount of epochs necessary to reach 93% accuracy. Data points and error bars represent the average and standard deviation, respectively (*n* = 20 runs with random initializations).

To further substantiate these findings, we investigated the networks’ responses to a standardized input at different sparsity levels. Analyses revealed that while the average firing rate of the untrained network decreased with increasing sparsity, the training process regularized the network’s overall activity, such that the average firing rate converged to a constant value independent of the connection density (Fig. 3A). This value was found to be approximately 16 Hz, which was determined by the balance between the classification loss minimization and a regularization term for the firing rate (see Supplementary Materials).

**Figure 3:**
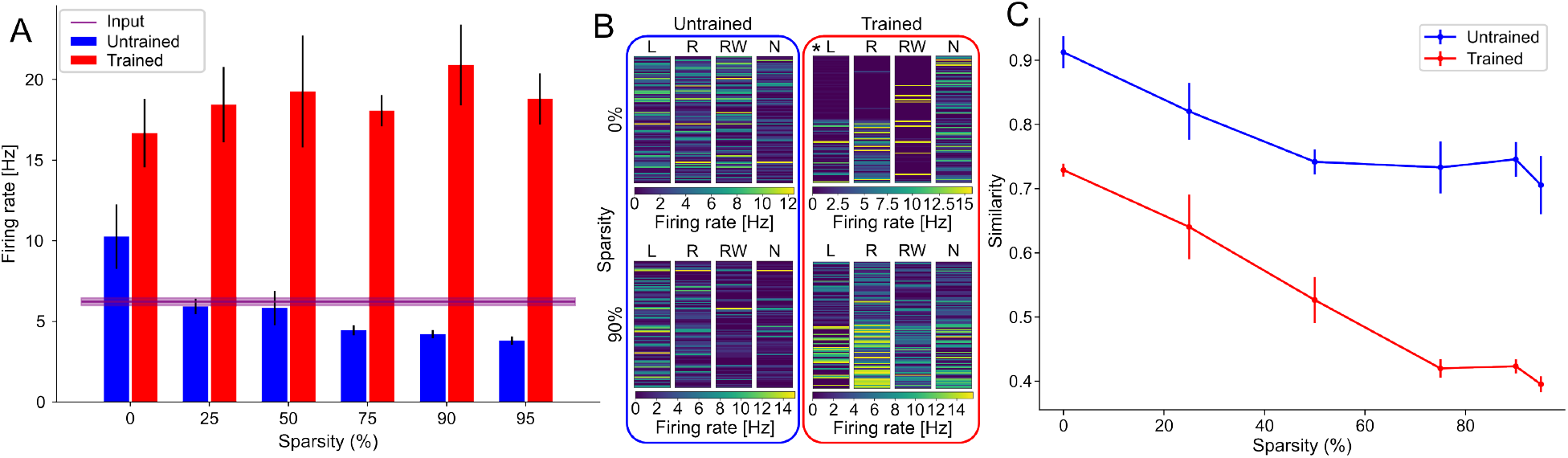
Impact of sparsity on network dynamics. A) Average firing rate during training for RSNNs with different sparsity. B) Firing rate of the recurrent-layer neurons during standardized episodes. RSNNs at 0% and 90% sparsity for untrained (left) and trained networks (right) are compared. Data are categorized by input type: “L” and “R” represent responses to the left and right signals, respectively, “RW” stands for the reward signal, and “N” represents the silence period when the input is only noise. * indicates that the training didn’t reach the accuracy threshold of 93% within the maximum amount of epochs (500). C) Cosine similarity between responses of the network to left and right signals, measured before and after training at different levels of sparsity. Panels A and C represent averages over 10 runs with error bars showing standard deviations.

The effect of training was also apparent on network responses to different input signals (Fig. 3B). Comparing the cosine similarity between the network’s responses to the left and right input channels, we observed a consistent reduction in similarity post-training (blue vs. red curves), indicating an enhanced capacity for input discrimination (Fig. 3C). Furthermore, the similarity decreased with increasing sparsity, suggesting that more sparsely connected networks are intrinsically more effective at differentiating between input signals. This ability to generate distinct activation patterns may underlie the improved performance observed at higher levels of sparsity.

### E-prop-trained input layer filters out noise

Next, we studied how different levels of input noise affected the performance of the e-prop algorithm. To control the level of spontaneous activity in RSNNs, we modulated the frequency of the Poisson process that generates input spikes in the noise channel. Some examples of the inputs employed during the experiment are reported in Fig. 4A. The input noise we employed in this experiment is asynchronous and uncorrelated, which resembles thalamic spontaneous activity [34]. Network sparsity was kept fixed at the optimal value obtained in the previous section, namely 92%.

**Figure 4:**
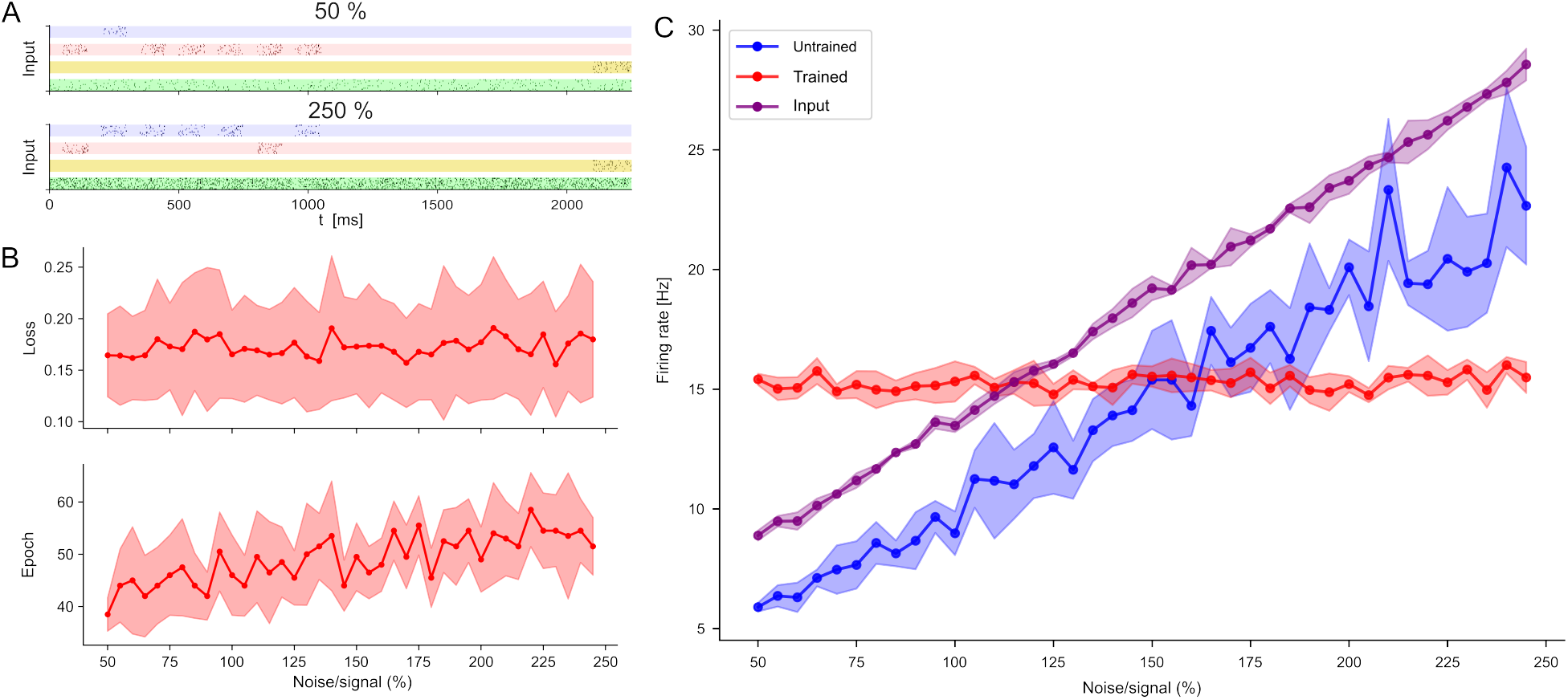
A) Examples of inputs at 50% (top) and 250% (bottom) noise-to-signal ratio (NSR). B) Top: average loss between epochs 400 and 500. Bottom: average number of epochs required to reach 93% accuracy. C) Average firing rate during training at different NSR. Firing rates are plotted in blue and red for the untrained and trained networks, respectively.

Fig. 4B shows the network’s change in behavior with the increase of noise. We see that the epochs required to reach the desired accuracy of 93% increased from an average of 38.5 ± 3.2 at 50% noise-to-signal ratio (NSR) to 51.5 ± 5.5 at 250% NSR (*n* = 20 samples; two-sided *t*-test, *p <* 0.01), however the change stayed within 2*σ* and the network was still capable of solving the task at very high noise levels. While the loss showed a modest change from 0.16 ± 0.04 at 50% NSR to 0.18 ± 0.06 at 250% NSR, this change was not statistically significant (two-sided *t*-test, *p >* 0.05).

To further characterize the network activity under various noise intensity, we evaluated the mean firing rate of the recurrent layer at different stages of training and analyzed its behavior in response to a standardized input with different amounts of noise. As summarized in Fig. 4C, the untrained network’s activity grows proportionally to the density of input spikes, while the trained network consistently maintained a firing rate of approximately 16 Hz, independent of the noise injected. This result extends the data from the previous section, showing that the network also converges towards a stable firing rate independently of the amount of noise injected. The ability of the network to solve the complex temporal credit assignment task, even under high NSR conditions, suggests an intrinsic mechanism for noise filtering in the input layer that stabilizes network activity in the recurrent layer.

### Network mechanisms of e-prop training

Finally, we examined the network’s evolution during training to better understand how the synaptic landscape is shaped by training. Analysis revealed that all three layers evolved through time and thus contributed to learning (Fig. 5A). We also verified that changes in the input and recurrent weight matrices, both trained under e-prop, plateaued at approximately 60 epochs around which the accuracy threshold of 93% was met (Fig. 5A, center). These results confirm that e-prop converged to an optimal state for solving the task at hand. While the norm of the readout weight matrix did not saturate within 500 epochs, perhaps reflecting the typical convergence characteristics of the standard gradient descent, it had only a mild impact on the accuracy (Fig. 5A, right).

**Figure 5:**
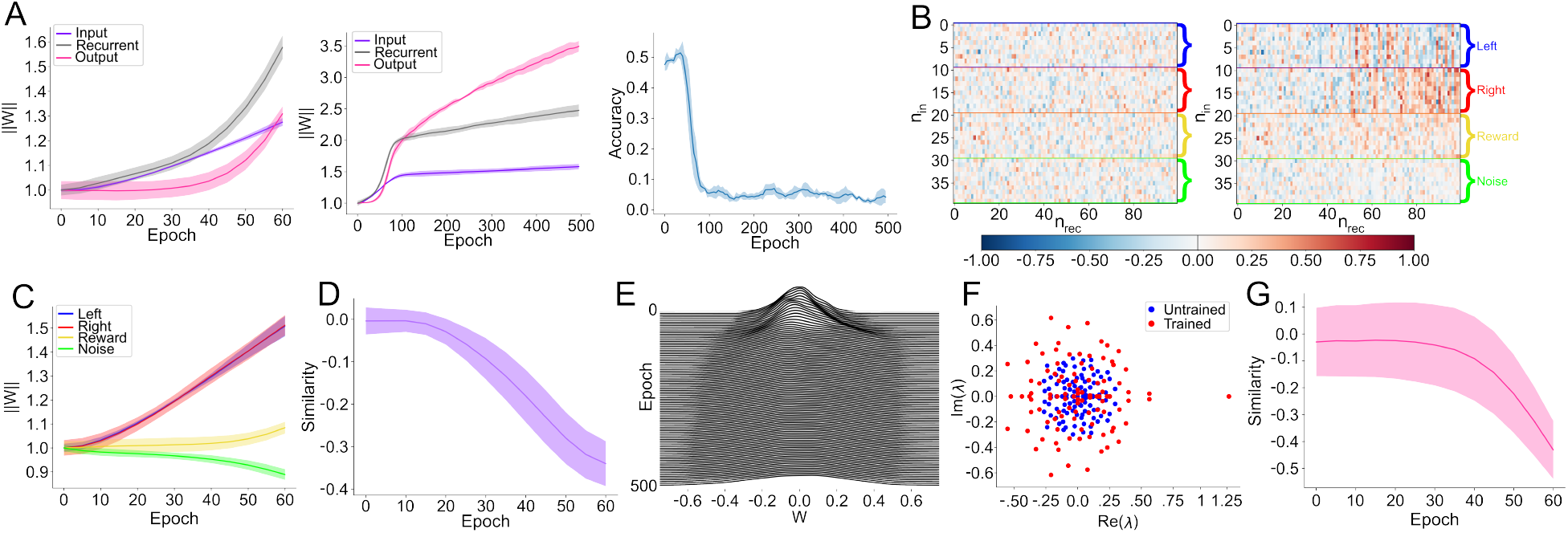
Evolution of network weights during training. A) Frobenius norm of the input, recurrent, and output weight matrices, plotted up to the point when the 93% accuracy threshold is reached (left) and over 500 epochs (center). The norms are normalized to their initial values to highlight weight growth. Right: Evolution of the accuracy during training. Note that the accuracy here was estimated from the training data, while the convergence criterion at the threshold accuracy of 93% used elsewhere was based on evaluation using the testing data. B) Input weight matrices before (left) and after training (right). C) Norm of each input channel throughout training. D) Cosine similarity between left and right input channels. E) Histogram of recurrent weights over the course of training (500 epochs). F) Eigenvalue spectra of the recurrent weight matrix for the untrained (blue) and trained (red) network. G) Cosine similarity between left and right output channels. Each curve in panels A, C, D, and G represents the average over 10 different runs, with the shaded error bars denoting the standard deviation. The results were obtained setting the network sparsity to 92% and the NSR to 250.

In the input layer, each input channel exhibited distinct patterns of evolution (Fig. 5B). In particular, as suggested in the previous section, the layer autonomously learned to filter noise and highlight input signals. As can be seen in Fig. 5C, the norm of the signal channels increased, while that of the noise channel was suppressed.

To gain further insight into how the left and right channels evolved during the training process, we computed the cosine similarity between their respective weights over time (Fig. 5D). Analysis revealed a pronounced decrease in similarity, indicating that the input layer evolved to route the two input channels to distinct neuron populations of the recurrent layer. This divergence is consistent with the distinct activation patterns observed in previous sections. Similar mechanisms have been observed in the sensory complex in the brain. In particular, the thalamus has been shown to adaptively filter inputs in response to feedback from the cortex [35] and to route information to bootstrap learning [36, 37]. These findings here explain the network’s resilience to injected noise and its enhanced performance at higher sparsity.

Analysis of the recurrent layer revealed a pronounced widening the weight distribution throughout training, while roughly maintaining the initial balance of excitation and inhibition (Fig.5E). Furthermore, the eigenvalue spectrum of the recurrent weight matrix broadened substantially, with the spectral radius increasing up to a value of 1.25 (Fig. 5). This behavior of the spectral radius is usually associated with a system approaching the “edge of chaos”, a regime of operation characterized by enhanced temporal stability and memory capacity [38, 39].

Lastly, we evaluated the evolution of the output layer. Consistent with the input layer, the two output channels evolved their synaptic weights to readout from different neurons in the recurrent layer. This differentiation is reflected in a decrease in cosine similarity between the left and right output channels (Fig. 5G), supporting our hypothesis that the network learns to route distinct neuron populations in the recurrent layer to each output channel.

## IV. DISCUSSION

In this study, we characterized the behavior of a RSNN trained using e-prop at different connection density and strengths of input noise. Simulations revealed that sparser networks are more effective at translating different inputs into distinct activation patterns in the recurrent layer which makes them better at solving the complex temporal credit assignment task. It was also found that the input layer adapts to act as a filter, making the network particularly resilient to noise.

Building on the biological plausibility claim of e-prop [22], we delved deeper into the learning mechanisms by demonstrating a thalamus-like functionality in the input layer and showing improved performance in networks with more biologically-plausible connectivity. The results of this paper reinforce the evidence found in [40] about the resemblance between the input layer and the thalamus in suppressing irrelevant input information such as noise. Our findings also extend previous work on the role of sparsity in ANNs [41] by demonstrating similar benefits in RSNNs trained with a biologically plausible algorithm. Additionally, the results confirm the earlier findings on the increased efficacy of sparser RSNNs in solving classification tasks [42], generalizing the conclusion to tasks with long temporal delays between inputs and rewards.

Results herein lay down a basis for optimization of RSNN morphology to solve classification tasks. Future works may explore more complicated network structures, such as the modular structure we explored in the past using biological neurons [43, 44]. It will also be interesting to explore other means to simulate spontaneous activity, background activity that occurs even in the absence of an input signal, and originates from various sources [45]. Although this property is typically disregarded in conventional artificial neural networks for machine learning, it constitutes a fundamental aspect of biological neural networks. There are multiple theories about the role this spontaneous activity plays in a plethora of neural processes, such as contributing to consolidation and retrieval in memory [46], modulating the integration capabilities of neurons in sensory tasks [47] or introducing variability to computation in cognitive and sensory applications, similarly to noise in machine learning [48, 49]. It would also be interesting to study the impact of connectivity and noise on unsupervised training and reinforcement learning, as well as applying the e-prop learning algorithm to a biologically inspired computational paradigm as in [50].

The features of the e-prop algorithm we highlighted are promising for its implementation in neuromorphic hardware: in particular, the intrinsic mechanism that minimizes the impact of noise and the sparse connectivity found to be optimal could significantly alleviate the hardware requirements of the implementation and simplify the design process for an ASIC such as ReckOn [25]. While further studies employing more diverse and complex datasets are needed to fully validate the generalizability of our conclusions to other tasks, prior work in RSNN suggests that the impact of network topology on performance is mostly task-independent [51, 52]. These studies would advance the application of biologically inspired approaches in neuromorphic systems, while simultaneously providing insights into the functional significance of biological architectures shaped by evolution.

## Acknowledgments

The work was partly supported by MEXT Grant-in-Aid for Transformative Research Areas (A) “Multicellular Neurobiocomputing” (24H02330, 24H02332), JSPS KAKENHI (22K19821, 22KK0177, 23H02805, 23K24913, 23K28179, 25H00447), JST-CREST (JPMJCR19K3), JST ALCA-Next (JPMJAN23F3), the WISE Program for AI Electronics by Tohoku University, and the Cooperative Research Project Program of the Research Institute of Electrical Communication (RIEC) at Tohoku University. This research was partly carried out at the Laboratory for Nanoelectronics and Spintronics, RIEC, Tohoku University.

## Data availability

The data that support the findings of this study are available upon reasonable request to the authors.

## Supplementary Materials

### Trace computation

In this section we will delve deeper into the computation of the traces for our particular experimental setup following the original paper from Bellec et al. [1]. For LIF neurons, the only value describing the internal state (**h**_*i*_) is the membrane potential (*v*_*i*_), thus the eligibility vector (*ε*_*ij*_) can be obtained by:

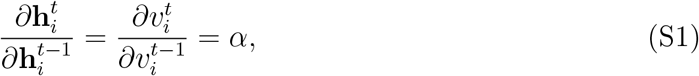

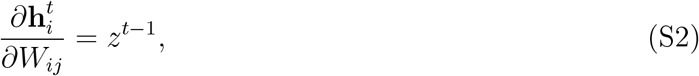

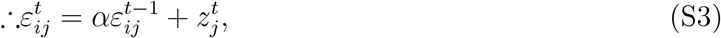

where *W*_*ij*_ is the weight matrix, and *α* is the decay constant of the LIF neurons. *ε*_*ij*_ is in fact a low-pass filtered version of the spike vector (*z*_*i*_) for the neuron. We define this low-pass filter for a quantity *q* as 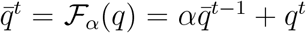. Thus, the eligibility trace (*e*_*ij*_) takes the form:

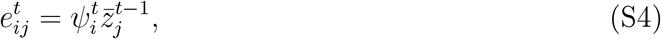

where 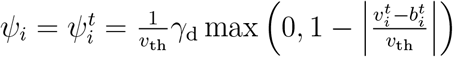.

For ALIF neurons, their internal state is represented by the vector 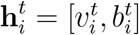, where *b*_*i*_ is the threshold adaptation, thus the eligibility vector takes the form 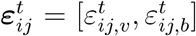 . The quantities in Eq. 9 of the main text are computed as:

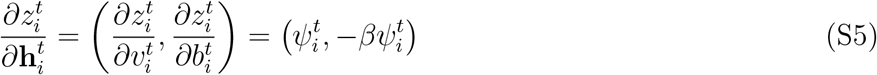

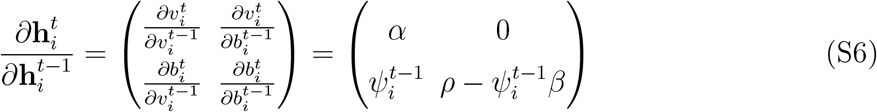

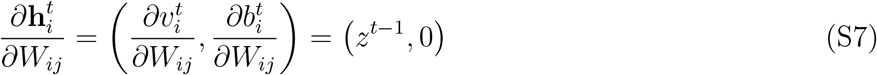

where *ρ* represents the decay constant for the adaptive threshold, and *β* is a constant. We can now compute the contribution given by the adaptable threshold to the eligibility vector:

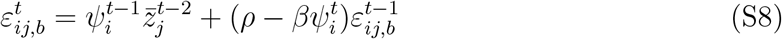

The final trace is thus given by:

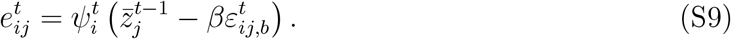

### Firing rate regularization

In this study, we introduced a regularization mechanism to ensure the stability of the firing rate during network activity. The regularization term *E*_reg_ is defined as:

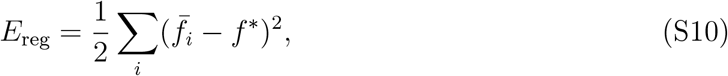

where 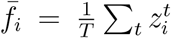 is the average firing rate of the *i*th neuron over a learning episode of duration *T*, and *f* ^∗^ is the target firing rate, set to 10 Hz. This additional loss term contributes to the evolution of the network in parallel to the e-prop rule described in the main text, and is incorporated into the total loss function *E* as:

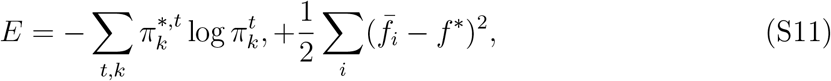

leading to the following plasticity rule:

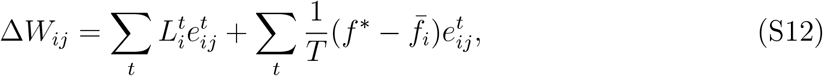

where *L*_*i*_ is the learning signal defined in the main text.

